# Bayesian inference of metabolic kinetics from genome-scale multiomics data

**DOI:** 10.1101/450163

**Authors:** Peter C. St. John, Jonathan Strutz, Linda J. Broadbelt, Keith E.J. Tyo, Yannick J. Bomble

**Affiliations:** Biosciences Center, National Renewable Energy Laboratory, 15013 Denver West Parkway, Golden CO 80401, USA; Department of Chemical and Biological Engineering, Northwestern University, 2145 Sheridan Road, Evanston IL 60208, USA

## Abstract

Modern biological tools generate a wealth of data on metabolite and protein concentrations that can be used to help inform new strain designs. However, integrating these data sources to generate predictions of steady-state metabolism typically requires a kinetic description of the enzymatic reactions that occur within a cell. Parameterizing these kinetic models from biological data can be computationally difficult, especially as the amount of data increases. Robust methods must also be able to quantify the uncertainty in model parameters as a function of the available data, which can be particularly computationally intensive. The field of Bayesian inference offers a wide range of methods for estimating distributions in parameter uncertainty. However, these techniques are poorly suited to kinetic metabolic modeling due to the complex kinetic rate laws typically employed and the resulting dynamic system that must be solved. In this paper, we employ linear-logarithmic kinetics to simplify the calculation of steady-state flux distributions and enable efficient sampling and variational inference methods. We demonstrate that detailed information on the posterior distribution of kinetic model parameters can be obtained efficiently at a variety of different problem scales, including large-scale kinetic models trained on multiomics datasets. These results allow modern Bayesian machine learning tools to be leveraged in understanding biological data and developing new, efficient strain designs.

## Introduction

Optimizing the metabolism of microorganisms for maximum yields and titers is a critical step in improving bioprocess economics (Davis et al., 2013). Towards this goal, characterization of engineered strains has become increasingly detailed with the growing availability of transcriptomic, proteomic, and metabolomic analysis techniques (Zhang et al., 2009). These methods, collectively termed multiomics, measure relative changes in gene, protein, or metabolite concentrations across different strains or growth conditions (Hackett et al., 2016). However, utilizing multiomics data to make informed decisions about future strain improvements remains a major challenge in modern bioengineering (Marcellin and Nielsen, 2018). Cellular metabolism is controlled by a vast network of enzymes with complex and nonlinear kinetics, while regulatory effects (both allosteric and transcriptional) add additional layers of complexity to these metabolic systems. While the field of systems biology has developed modeling frameworks that can describe these types of interactions in great detail, parameterizing these models from indirect, *in vivo* data is challenging and typically infeasible at the genome-scale (Saa and Nielsen, 2017). There is therefore a need for mechanistic modeling frameworks that can handle the large amounts of data generated through multiomics experiments to yield actionable insight for strain engineers.

Traditionally, multiomics data have been understood through statistical approaches by searching for genes, proteins, and metabolites whose activity levels are correlated with improved product production (Yoshikawa et al., 2012). While useful in identifying potential targets, statistical methods that examine multiomics data without consideration for their interconnected nature may miss important trends. Conversely, metabolic modeling frameworks that readily reconcile connections between metabolites, fluxes, and proteins can have difficulty in using multiomics data to improve their predictions. Stoichiometric models of metabolic networks have proved among the most successful techniques for incorporating existing knowledge of genomic and biochemical networks into strain designs (Orth et al., 2010). Instead of attempting to estimate the parameters of detailed rate rules for each enzymatic reaction in the cell, constraint-based models assume that metabolic reactions reach a pseudo-steady state with respect to the longer time scales of cell growth and substrate depletion. These methods there investigate feasible steady-state phenomena in a parameter-free approach by placing constraints on reaction fluxes in accordance with stoichiometric (Orth et al., 2010), thermodynamic (Henry et al., 2007), enzymatic (Sánchez et al., 2017), and regulatory (reviewed in (Saha et al., 2014)) rules. The resulting models are invaluable for predicting ways to restrict cell physiology to specific regions (e.g. forcing growth-coupled product production through gene knockouts (Burgard et al., 2003)). However, their parameter-free construction renders the direct inclusion of measured metabolite and enzyme concentrations difficult, and their lack of kinetic information makes them poorly suited to recommending enzyme expression changes to optimize pathway flux. Additionally, constraint-based techniques typically assume growth as a cellular objective, making them poorly suited to *in vitro* or other non-growth associated bioprocesses.

While some studies have used constraint-based techniques to interpret multiomics data (Brunk et al., 2016; Cotten and Reed, 2013; O’Brien et al., 2014; Sánchez et al., 2017; Yizhak et al., 2010), building and parameterizing kinetic models will likely be essential in utilizing these types of data to predict metabolic interventions that will improve flux through a given pathway. Kinetic models of metabolism typically describe the interior cellular environment through systems of coupled, nonlinear ordinary differential equations with metabolite concentrations as the state variables. Metabolite concentrations change with time according to the kinetics of enzyme-mediated reactions, reaching a stable steady-state for constant concentrations of extracellular substrates. Understanding systems-level effects is the main motivation of metabolic control analysis (MCA), which links effects of local perturbations (i.e., changes to enzyme expression) to changes in the resulting steady-state concentrations and fluxes (Ehlde and Zacchi, 1997; Visser and Heijnen, 2002). MCA defines local coefficients, or *elasticities*, which are local approximations to the reaction rate rule and relate reaction flux to substrate concentration. Through linearization, MCA also solves for global *control coefficients*, which relate steady-state fluxes and concentrations to enzyme levels. Parameters estimated via MCA are directly relevant to strain engineering goals, as they allow the prediction of which enzymes to over- or underexpress to achieve a desired change in pathway flux. As a result, a number of computational frameworks have been developed to perform MCA with incomplete data (Chakrabarti et al., 2013; Delgado and Liao, 1991; Wang et al., 2004; Wu et al., 2004). Most MCA approaches require an accurate dynamic model of metabolism that must be informed from experimental measurements. However, direct measurements of enzyme kinetics in vivo are difficult, and measurements *in vitro* do not always accurately reflect in vivo dynamic behavior (Teusink et al., 2000; Zotter et al., 2017). Experimental characterization of genome-scale enzyme kinetics is particularly challenging (Nilsson et al., 2017). As a result, the most readily available data for parameterizing kinetic metabolic models is obtained by repeatedly perturbing enzyme expression or external metabolite concentration and characterizing the resulting strain’s pseudo-steady-state behavior through multiomics experiments.

The process for constructing a kinetic metabolic model therefore consists of choosing an appropriate functional form for the rate rule of each reaction and estimating parameter values from experimental data. A number of different frameworks for describing the kinetics of enzyme-catalyzed reactions have been proposed, covering a spectrum of computational efficiency and mechanistic fidelity (reviewed in (Heijnen, 2005; Saa and Nielsen, 2017)). However, regardless of the framework chosen, estimating uncertainty in fitted parameter values is a challenging process due to the dimensionality of the resulting parameter space. One approach is to find an ensemble of possible parameter values that when passed through the kinetic model closely reproduce the observed experimental data (Tran et al., 2008). These distributions in parameter values can then be updated as more data is collected and used to predict enzyme targets that give the most likely chance of improving performance (Contador et al., 2009). This technique has more recently been formalized as Bayesian inference (Saa and Nielsen, 2016), where parameter values are modeled as probability distributions. In Bayesian inference, a *likelihood model*, *p*(*y*|*θ*), is constructed for the probability of observing the measured data, *y*, given particular values for each parameter, *θ*. Combined with a *prior* distribution, *p*(*θ*), for each parameter that represents generally feasible values, numerical approaches use Bayes theorem,

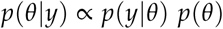

to estimate the posterior parameter distribution *p*(*θ*|*y*): the probability a parameter takes the given value after accounting for the observed data.

A major limitation of ensemble-based modeling has been its ability to scale both to larger datasets as well as larger kinetic models. Computation times for metabolic ensemble modeling (MEM) on the order of hours per parameter sample have been encountered even for relatively small models and datasets in previous studies (Saa and Nielsen, 2016; Tran et al., 2008). Despite improvements to the computational efficiency of MEM (Greene et al., 2017; Zomorrodi et al., 2013), applications of ensemble modeling have been restricted to small datasets and/or kinetic models, without the ability to scale to genome-scale kinetic representations fit with data from multiomics workflows. This limitation stems from the need to perform computationally intensive integration of underlying ODEs to find steady-state concentrations and fluxes, combined with a relatively inefficient rejection sampler approach used to estimate the posterior distribution. Modern and efficient inference algorithms (Hoffman and Gelman, 2014; Kucukelbir et al., 2017) require information on the gradient of the likelihood function with respect to the kinetic parameters, which can only be obtained for ODE models through numerically intensive ODE sensitivity analysis (Li and Petzold, 2000).

In this study, we present a scalable method for inferring posterior distributions in kinetic parameters of large metabolic models with multiomics datasets. We sidestep many of the previously discussed computational bottlenecks through the use of linear logarithmic (linlog) kinetics as an approximate reaction rate rule (Visser and Heijnen, 2003; Visser et al., 2004). Linlog kinetics is derived using the thermodynamic concept that reaction rate is proportional to reaction affinity near equilibrium (Onsager, 1931). While many biochemical reactions are far from equilibrium, this relationship remains linear over a wide range of reaction affinities for enzymatic reactions (Meer et al., 1980; Rottenberg, 1973), As an approximation, linlog kinetics does not describe enzyme-mediated kinetics as faithfully as more mechanistic frameworks (Saa and Nielsen, 2017). However, linlog kinetics has been shown to be accurate up to 20-fold changes in metabolite concentrations (Visser and Heijnen, 2003), and for 4 to 6-fold changes in enzyme concentration relative to a reference state (Visser et al., 2004). As a result, linlog kinetics has been used as a framework for estimating flux control coefficients from a range of data sources (Heijnen et al., 2004; Kresnowati et al., 2005; Nikerel et al., 2006, 2009). Most importantly, this kinetic formalism allows steady-state fluxes and metabolite concentrations as a function of enzyme expression to be determined directly via linear algebra, without the need to explicitly integrate the dynamic system until a steady-state is reached (Visser et al., 2004). We are therefore able to leverage modern Bayesian inference and machine learning algorithms, including Hamiltonian Monte Carlo (HMC) (Neal, 2010) and variational inference (Blei et al., 2017) to fully characterize the posterior space. Additionally, this framework naturally lends itself to directly incorporating relative changes in metabolite and protein concentrations between experimental conditions, without requiring absolute quantification.

We show that this method is capable of providing systems-level insight into metabolic kinetics through estimated distributions in control coefficients for a wide range of kinetic model and dataset sizes. First, we demonstrate the method on a simple *in vitro* example, showing that the method is flexible enough to capture allosteric interactions between metabolites and enzymes. We next show that the method appropriately captures uncertainy in estimated parameters, revealing significant flux control coefficients for only the most likely enzyme perturbations in the case of limited biological data. Finally, we employ the method to integrate thousands of individual metabolomic, proteomic, and fluxomic data-points with a large-scale model of yeast metabolism. We therefore show that the field of metabolic modeling can take full advantage of recent advances in the fields of probabilistic programming, machine learning, and computational statistics, and that ensemble-based approximate kinetic modeling approaches are capable of scaling to genome-sized models and datasets to provide interpretable and actionable insight for strain engineers.

## Results and discussion

### Enabling efficient Bayesian inference through linlog kinetics

We begin with a review of the relevant equations from dynamic flux balance analysis and the linear-logarithmic kinetic framework, which together form the theoretical basis for the methodology discussed in the remainder of the study. In flux balance analysis, we assume that metabolite concentrations, *x*, quickly reach a pseudo-steady state by balancing fluxes *v* through each reaction.

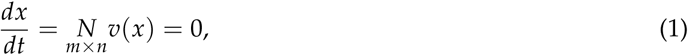

for *n* reactions and *m* metabolites, where *N_ij_* indicates the stoichiometry of metabolite *i* in reaction *j*. Linlog kinetics approximates a reaction rate *v*(*x*) as a sum of logarithms (Visser and Heijnen, 2003). For the reaction *A* → *B* + *C*, the reaction rate is modeled as

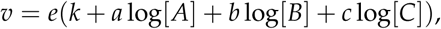

for which the coefficients *a* > 0; *b,c* < 0 allow an approximation of Michaelis-Menten-type kinetics (Figure 1). This approximation is most accurate in the vicinity of an introduced reference state, *e*^∗^, *v*^∗^, *x*^∗^ (Visser and Heijnen, 2003). As the goal of the proposed method is to tailor enzyme expression to maximize desired fluxes, the reference state is best chosen as the current optimal performing strain. Deviations from this state can be described by the flux expression

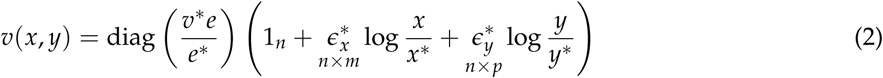

where *y* is the concentration of *p* external (independently controllable) metabolite species, and 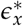 and 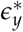 are sparse matrices of kinetic parameters describing the effects of changes to metabolite concentrations on reaction rates. Elasticities parameterize the slope of the reaction rate rule near the reference state. Linear-logarithmic kinetics therefore offer a close approximation to standard Michaelis-Menten kinetics in the vicinity of the reference state concentration (*x*^∗^). A benefit of the linlog approximation is that enzyme elasticities are direct kinetic parameters. Since these slopes tend to be positive for reactants, negative for products, and not be much larger than 1, reasonable starting guesses and bounds can be generated for all kinetic parameters in the model in a much easier fashion and for rate rules parameterized through traditional enzymatic expressions. Elasticities for linear logarithmic kinetics have typically been estimated in the literature using multiple linear regression (Chen et al., 2017; Wu et al., 2004), where estimated fluxes for each reaction are fitted as a function of their measured metabolite concentrations. However, this approach does not enforce the *Nv* = 0 constraint, nor does it allow for missing data in concentration or flux measurements. We demonstrate that incorporating steady-state constraints is computationally feasible, and that a full characterization of the posterior space can be accomplished using Hamiltonian Monte Carlo.

While linlog kinetics is a close approximation of more mechanistic rate rules, it suffers from a number of notable inconsistencies. One consequence is that fluxes can approach negative infinity as metabolite concentrations approach zero, making the framework unsuitable for describing complete pathway knockouts. However, in practice metabolite concentrations are typically expressed as log-transformed variables which also cannot fall to zero. Other methodological strategies discussed later also prevent fluxes from taking unrealistic values, including using a least-norm linear solve for steady-state concentrations and clipping data to a finite range. Additionally, as a local approximation, the method will poorly reproduce alternative rate rules at large deviations from the reference state. However, because cellular systems are constrained by homeostasis, metabolite concentrations generally do not change drastically enough to invalidate rate estimates (Ishii et al., 2007).

A key step in dynamic modeling of metabolic networks is solving for the steady-state concentrations and fluxes that arise from a given parameterization. Simulating this perturbation efficiently with the mathematical model is therefore a key step in estimating parameter values for the *ε*^∗^ matrices. In doing so, it is useful to define transformed variables in order to rewrite Equation 2 in a linear form (as demonstrated by Smallbone et al., 2007):

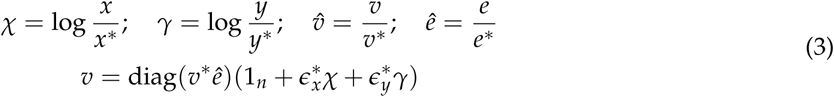

Since log-transformed metabolite concentrations are linearly related to the reaction fluxes, concentrations which yield steady-state behavior can therefore be determined via a linear solve (Visser and Heijnen, 2003) after combining Equation 3 with Equation 1:

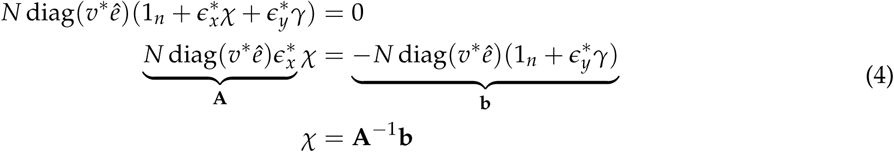

This significant result is the key advantage of linlog kinetics over alternative nonlinear rate laws. While determination of steady-state concentrations would typically require a computationally intensive ODE integration, in this approximation they can instead be calculated using a single linear solve. Additionally, it is relatively straightforward to obtain forward and reverse-mode gradients for this operation (changes in steady-state with respect to changes in kinetic parameters), a much more difficult task for ODE integration (Petersen and Pedersen, 2012).

However, in general a metabolic system will contain *conserved moieties,* or metabolite quantities which can be expressed as linear combinations of other metabolites (*e.g.* ATP + ADP = constant). The stoichiometric matrix *N,* and as a result the A matrix defined above, will therefore not be full row rank. In effect, this means that Equation 4 has multiple solutions, each of which corresponds to a different total cofactor pool. In metabolic control theory, this problem has traditionally been solved through the introduction of a link matrix, L, and a reduced set of metabolites with conserved moieties removed (Smallbone et al., 2007; Visser and Heijnen, 2002). Through the link matrix, the matrix A can be transformed to a full-rank, square matrix and a unique steady-state can be determined that corresponds to the dynamic system’s true steady-state. However, in most biological experiments, changes to steady-state enzyme expression correspond with separately cultured cell lines for which the assumption that total cofactor pools would remain constant is not necessarily valid. Instead, we propose that a more biologically relevant solution to Equation 4 is one that minimizes ∥*χ*∥_2_: *i.e.*, the solution that results in the smallest deviation of metabolite concentrations from the reference state. This assumption has experimental support in that intracellular metabolite concentrations tend to be buffered from drastic changes through feedback circuits at the genetic and enzyme level (Ishii et al., 2007). We therefore calculate steady-state metabolite concentrations through a pseudoinverse,

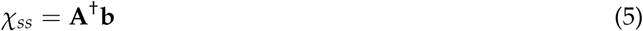

A derivation of the forward and reverse-mode gradients for the regularized linear solve operation is included in the supplemental text. We note that in practice, numerical stability is improved if **A** can be made full row-rank prior to the least-norm linear solve. We can therefore replace *N* with 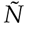 (by removing rows corresponding to redundant conservation relations) in order to form a wide *A* matrix (with more columns than rows) prior to performing the least-norm linear solve in Equation 5. Since a flux vector that satisfies 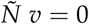 will also satisfy *N v* = 0, this change can be made without affecting the final solution.

Due to the changes to traditional MCA theory introduced by the altered steady-state calculation defined above, we also slightly modify the traditional calculations of metabolite and flux control coefficients (FCCs).

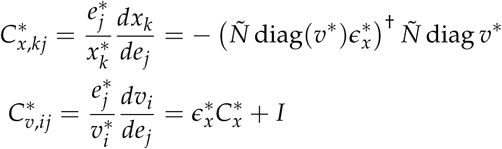

Since flux and metabolite control coefficient matrices describe the response of the steady-state to changes in enzyme expression, our altered versions describe the flux response at the particular steady-state in which metabolite concentrations are as close as possible to the unperturbed state. In practice, this has the effect of improving the identifiability of FCCs in the numerical experiments described below. A plot of FCC values obtained via both traditional and modified methods for the following genome-scale model is shown in Fig. S1, indicating that either both methods tend to yield a similar result, or that the identifiability of the link-matrix FCC is particularly poor, with the pseudoinverse FCC pulled close to zero.

With a suitable kinetic framework for calculating steady-state fluxes and concentrations as a function of enzyme expression, we next discuss the prior distributions and likelihood function required for Bayesian inference. The prior distributions represent our belief of possible parameter values before any experimental data is collected. For metabolite elasticity matrices we assume that for any given reaction, reactants are likely to be associated with a positive elasticity, while products likely have a negative elasticity (increasing reactant concentration increases reaction rate, while increasing product concentration decreases reaction rate). Alternatively, we assume that if a metabolite does not directly participate in a reaction, it can only regulate the reaction if it appears in the same sub-cellular compartment. We denote the vectors *c_m_* and *c_r_* of metabolite and reaction compartments, respectively. Since regulation of enzymatic reactions by otherwise nonparticipatory metabolites is relatively rare, we place a sparsity-inducing prior on its elasticity value. This distribution encourages elasticities for off-target metabolites to take values near zero, unless strong experimental evidence for a regulator interaction is present. The combined priors for enzyme elasticities can then be expressed through the following functional form, also depicted graphically in Figure 1.

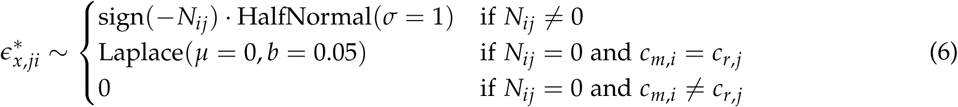

We note that the assumption that reactants and products must take positive and negative elasticity values, respectively, can be relaxed by replacing the half-normal distribution in Equation 6 with a skew-normal distribution with a positive shape parameter. This distribution reflects the belief that while reactants typically take positive elasticities, rare cases may exist where substrate inhibition results in a negative slope of reaction rate with respect to substrate concentration. In practice, however, this choice of a prior distribution results in less robust convergence to a stable posterior distribution and was avoided in higher-dimensional inference problems.

**Figure 1:**
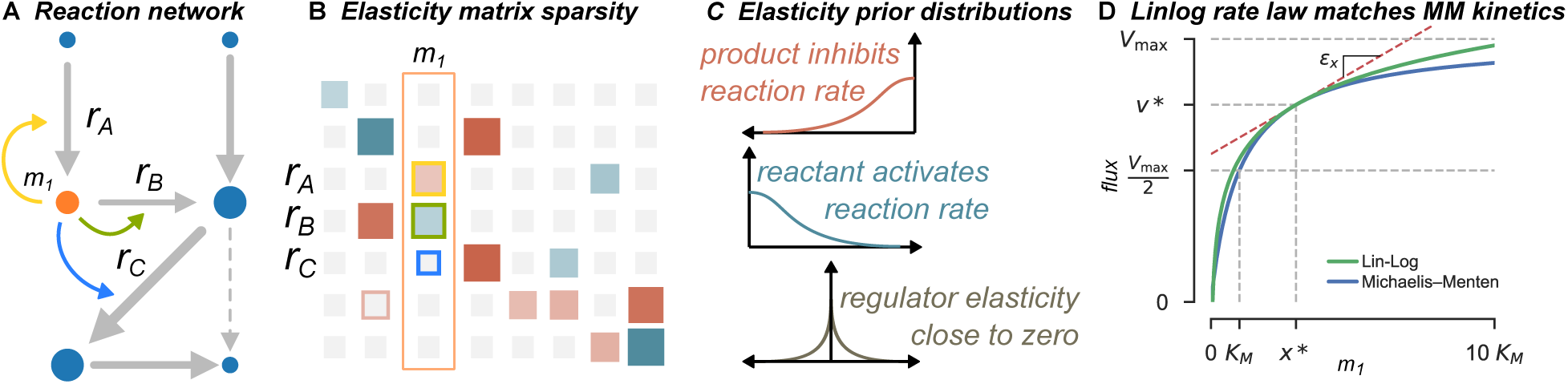
Overview of the modeling framework. (A) The stoichiometry of the reaction network is used to determine prior distributions for the elasticity parameters (B), represented in matrix form. (C) For a metabolite *m*_1_, the prior distribution has predominatly negative support for reactions in which the metabolite is a product (*r*_A_), positive support for reactions in which the metabolite is a reactant (*r*_B_), and a zero-centered, sparsity inducing prior for reactions in which the metabolite does not participate (*r*_C_). (D) The resulting rate law for linlog kinetics closely approximates Michaelis-Menten kinetics in the vacinity of the reference state.

An explicit likelihood function can be formed by constructing a statistical model for the observed data. We assume that observed data are normally distributed around the calculated steady-state metabolite and flux values.

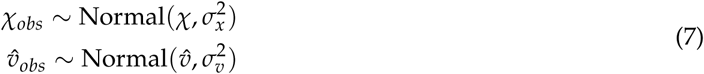

Experimental errors, *σ_x_* and *σ_v_*, can either be set explicitly or estimated from the data. For smaller-scale examples, we place half-normal priors on these variables, while for larger datasets we set these values explicitly to improve numerical stability. We also note that for genome-scale multiomics data, computational stability can be improved by fitting log-transformed normalized fluxes,

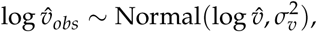

so that flux, metabolite, and enzyme expression data fall on similar orders of magnitude. While this assumption comes at the cost of preventing measured fluxes from reversing directions between perturbed states, this restriction was not significant for the examples considered in this study. However, this framework could be easily extended to handle situations where a measured flux reverses directions between experimental conditions. Most simply, the reversible reaction could be withheld from the log transform and fit in linear space. Alternatively, if separate estimates for the forward and reverse flux could be obtained, as is often the case in ^13^C labeling studies, the reaction could be decomposed and modeled separately as irreversible forward and reverse reactions.

Once the prior distribution and likelihood model have been specified, the remaining task is to numerically estimate posterior distributions in elasticity parameters Towards this goal, two inference algorithms were used. The No-U-turn sampler (NUTS) (Hoffman and Gelman, 2014), as a variant of Hamiltonian Monte Carlo (HMC), constructs an iterative process (a Markov chain) that eventually converges to the true posterior distribution. Markov chain Monte Carlo methods, while accurate, are computationally intensive and likely limited in application to smaller-scale models and datasets. While the major computational bottleneck in metabolic ensemble modeling (integrating an ODE until steady state) has been removed, calculating the likelihood function still involves a separate linear solve for each steady-state experimental condition. Therefore as model sizes approach the genome-scale, HMC methods quickly become computationally infeasible. Variational methods, however, offer an alternative to Markov chain Monte Carlo methods that can scale to models with thousands of parameters. Automatic differentiation variational inference (ADVI)approximates the posterior distribution by a simple, closed-form probability (typically Gaussian), then estimates parameters for the approximate posterior to minimize the distance between the true and approximated distribution.

### Characterization of an *in vitro* linear pathway

While the primary purpose of the proposed modeling framework is to parameterize genome-scale kinetic models from large, multiomics datasets, we first demonstrate the method on a simple pathway. We re-fit a simple three-reaction model (Wu et al., 2004) to steady-state *in vitro* flux and concentration data for a reconstructed subsection of lower glycolysis (Giersch, 1995). A schematic of the considered pathway is shown in Figure 2A. The model consists of two internal metabolite species, 2-phosphoglycerate (2PG) and phosphoenolpyruvate (PEP), and two metabolites with externally-controllable concentrations, adenosine diphosphate (ADP) and 2,3-bisphosphoglycerate (BPG). The model consists of three reactions in series, phosphoglycerate mutase (PGM), enolase (ENO), and pyruvate kinase (PK); therefore each carries the same flux at steady-state. The dataset consists of 19 separate experiments, each of which contains the enzyme loadings (concentrations) and external metabolite concentrations together with the resulting internal metabolite concentrations and steady-state flux.

Since all metabolites (including external species) are present in the same compartment, all elasticities are allowed to have allosteric interactions normalized with Laplace priors. Measurement errors in fluxes and metabolite concentrations were fit by the inference algorithm by placing a half-normal prior distribution on the *σ* values in Equation 7. The same reference steady-state was chosen (experiment 2) as was done by Wu et al., 2004.

Using NUTS, stable traces were found across four independent chains, indicating that each trace converged to the true posterior distribution (Figure 2B, Figure 5). For this small-scale example, NUTS sampling took less than 10 minutes on a single computer. Applying ADVI to this example, the evidence lower bound (ELBO), a measure of the closeness of fit between the approximated and true posterior distribution, converged after approximately 10,000 iterations of stochastic gradient descent (Figure 6). A full 25,000 iterations were completed in under 40 seconds on a single computer.

Comparing the results of the two inference methods indicates that both methods yield similar conclusions. ADVI fits a mean-field approximation - i.e., each parameter’s posterior is represented by a mean and standard deviation. Comparing the mean and variance of the elasticity posterior distributions from the two different approaches, we notice that while the mean values agree closely, ADVI underestimates the variance for many parameters (Figure 7). This underestimation is typical of mean-field ADVI (Kucukelbir et al., 2017), and might be alleviated in the future through more advanced variational methods (Rezende and Mohamed, 2015). A posterior predictive check for both inference methods indicates that the measured experimental data is well-captured by the model (Figure 2C). Despite normalizing priors on off-target regulation, all elasticity values in the internal metabolite elasticity matrix, 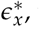 were confidently nonzero (as determined by whether the 95% highest posterior density (HPD) interval overlapped zero). Inferred regulatory interactions, which were all consistent between both inference methods, are shown in gray in Figure 2A. These include a strong repression of PGM by PEP, and a weaker repression of PK by 2PG. These off-target regulatory interactions (with similar elasticity values) were also found through the original linear regression approach of Wu *et al.,* 2004. For the external metabolite species, only one of the four possible off-target regulatory interactions, ADP activation of PGM, resulted in a posterior distribution that was confidently nonzero. This relatively weak interaction was rejected by the original linear regression method through a combination of experimental and mathematical reasoning, but underscores that interactions between metabolites and fluxes are inherently difficult to predict from this type of data: direct vs. indirect interactions often look similar, and causality is often impossible to establish. Notably, the posterior distribution as estimated via NUTS contains a rich amount of information on the identifiability of elasticity values (Figure 2D). Strong correlations in estimated parameters typically occur where the two elasticities share either a metabolite or reaction.

The main goal of the method is determining posterior predictive distributions in flux control coefficients, *i.e.*, determining major control points that determine how flux is distributed in the pathway. In this example, since steady-state fluxes are constrained to be equal for all three reactions, the FCCs are a vector of three coefficients that determine whether increasing enzyme concentration will increase or decrease pathway flux. Figure 2E shows posterior distributions in FCCs as estimated with both inference methods. These are compared against FCC distributions resulting from only the prior distributions on elasticity parameters, without considering any experimental results. Prior distributions are similar between all three enzymes and indicate no structural bias on flux control values. The data therefore indicate that pyruvate kinase (PK) is the limiting enzyme at the reference state. We also compare our FCC estimates against those originally calculated via linear regression, assuming specific allosteric interactions between metabolites and enzymes that differ from those found to be significant through our approach. Our estimates of flux control coefficients closely match those found by Wu *et al.*, 2004, indicating that systems-level properties are relatively insensitive to the particular parameterization of allosteric regulation.

The close agreement of the estimates provided by the approximate ADVI method to the more accurate NUTS sampling in elasticities and flux control coefficients is an important result. As most applications in metabolism involve a larger reaction network, approximate inference methods are likely the only techniques that will scale to biologically-relevant *in vivo* examples. We therefore rely only on these variational techniques for subsequent examples that deal with larger reaction networks.

**Figure 2:**
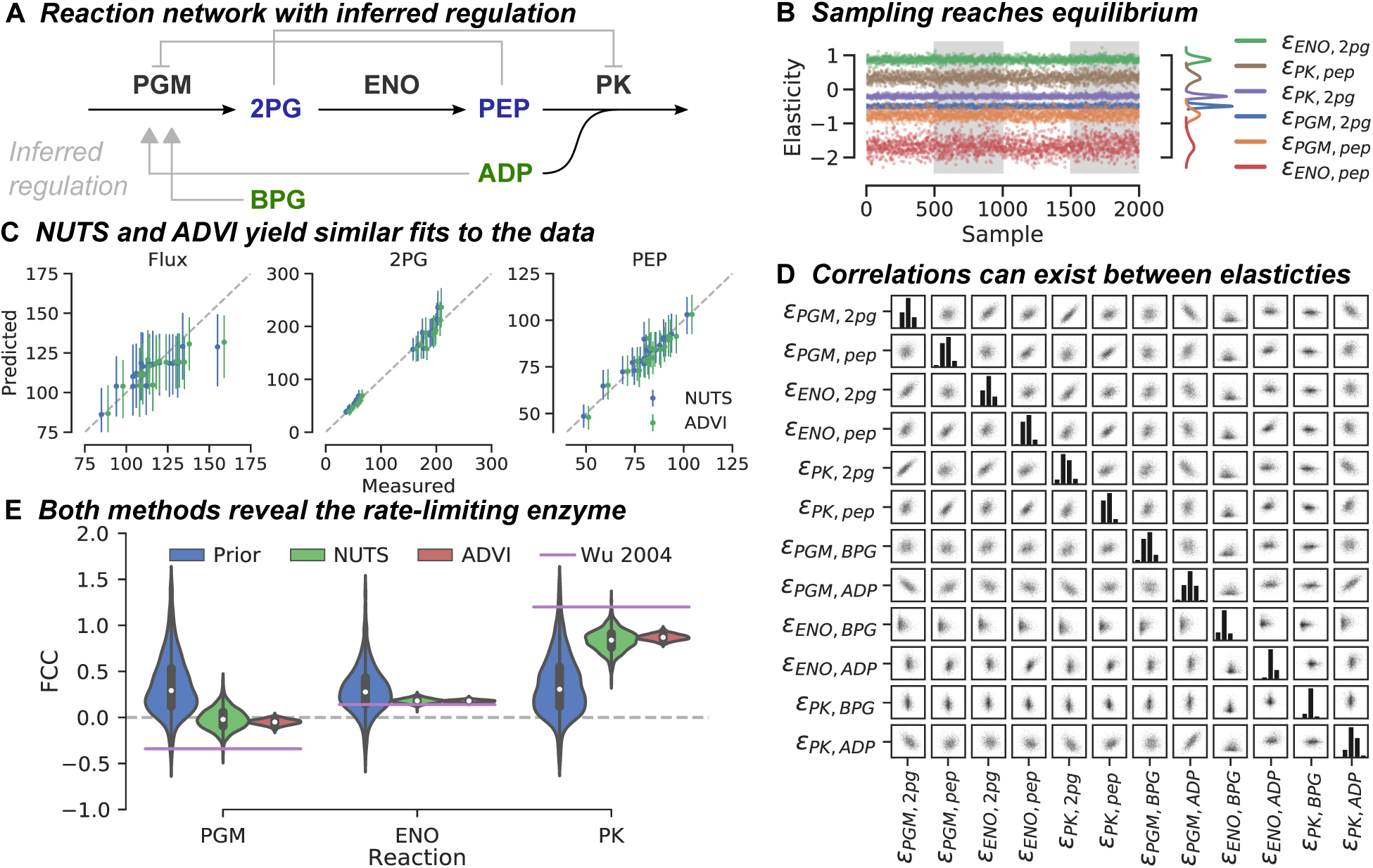
*in vitro* pathway inference. (A) Schematic of the considered pathway. Inferred allosteric interactions are shown in gray, in which arrows indicate an activation, while barheaded lines indicate inhibition. (B) Traces for 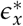 values as estimated by NUTS. Samples come from four parallel chains stacked together as indicated by the shaded regions. Resulting posterior densities are indicated by the inset on the right. (C) Posterior predictive distributions of steady-state flux and metabolite concentrations. Points represent medians of the posterior predictive distributions, with lines extending to cover the 95% HPD interval. Slight jitter was added to differentiate the distributions as estimated by NUTS and ADVI. (D) Pairplot of the posterior distributions of elasticity variables as estimated via NUTS. Strong correlations can exist between fitted parameters, which are missed by the mean-field ADVI approximation. (E) Violin plot of distributions in posterior flux control coefficients. Median and inner quartile range are indicated by the inner box plots, overlaid on a kernel density plot of each distribution.

### Determining optimal enzyme targets from limited data

We next demonstrate how the inference framework can be used to suggest enzyme targets in a many-reaction network, including branched reaction networks and conserved metabolite pools. The problem we consider was previously examined through ensemble metabolic modeling (Contador et al., 2009), and involves predicting what manipulations might further increase lysine production in engineered *E. coli* strains. We therefore replicate the previous ensemble modeling assumptions as closely as possible in order to allow a direct comparison of resulting predictions. The experimental data consists of six sequential enzyme overexpression experiments, all of which were observed to improve l-lysine yields (Kojima et al., 1993). The metabolic model used for inference comprises 44 reactions and 44 metabolites covering central carbon metabolism and lysine production, taken from Contador et al., 2009. A schematic of the reaction network is shown in Figure 3A.

As the goal of the inference approach is to estimate targets for subsequent lysine flux improvement, we chose the reference state for linlog kinetics to be the final, optimized strain with 5 overexpressed enzymes. Following the assumptions made in (Contador et al., 2009), we also assumed each overexpression doubled the concentration of the respective enzyme. Since the reference state was chosen to be the final, optimized strain, perturbed strains had lower relative enzyme concentrations and lysine flux. Reference state fluxes were also taken from previously published values, corresponding to a total lysine yield of 11.2% (Contador et al., 2009).

When analyzed with metabolic ensemble modeling, each successive enzyme overexpression was required to increase lysine flux over the previous base strain. However in our framework, we require a continuous and differentiable likelihood function. We therefore assume that each enzyme overexpression increases lysine flux relative to the wild-type strain by an additional 20% on average (with standard deviation 0.5%). Target relative fluxes after normalizing to the new reference are shown in Table 1. Prior distributions in enzyme elasticities were specified as described in Equation 6, and since the dataset did not include changes in external metabolites, no e_y_ values were needed. Posterior distributions were estimated using ADVI, with the optimization taking under three minutes. The posterior predictive distribution for each strain closely matches the target lysine fluxes, indicating the model is capable of reproducing the desired behavior (Figure 3B).

**Table 1:**
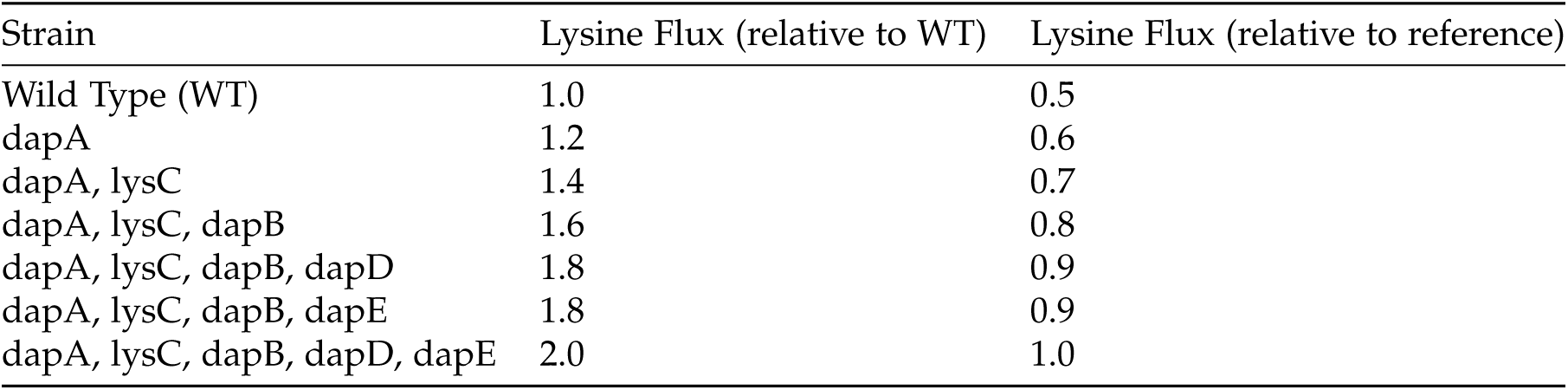
Assumed relative lysine fluxes for each considered strain, relative to both the wild-type flux and the chosen reference strain (dapA, lysC, dapB, dapD, dapE). Strain designs taken from (Kojima et al., 1993).

Using a half-normal distribution with *σ* = 1, prior distributions in elasticities associated with stoichiometric metabolite-reaction pairs had a 95% HPD that spanned from 0 to 2. Of these 133 ‘kinetic’ elasticity terms, only twelve were constrained by the experimental data to a 95% highest posterior density that spanned less than 0.75 elasticity units. In addition to these kinetic terms, three regulatory elasticities were identified as confidently nonzero (with a 95% HPD that did not include 0). These regulations include both feedback and feedforward connections, likely used by the model to fine tune the lysine expression to the desired 20% target in response to doubling of enzyme concentration. Posterior distributions for these elasticities are shown in Figure 3C, and confidently inferred regulatory interactions are shown in gray in Figure 3A. Unsurprisingly, nearly all of these elasticities involve reactions and metabolites in the lysine synthesis pathway, the only portion of the model for which overexpression results were provided.

Prior and posterior distributions in flux control coefficients were also calculated. Because only a limited selection of data was available to constrain the elasticity values, only five of the 44 reactions had a flux control coefficient for lysine export whose 95% HPD did not overlap zero. However, these five reactions were the same set of dapA, lysC, dapB, dapD, and dapE previously specified as successful modifications for improving lysine flux. Prior and posterior distributions in FCC values for lysine export are shown in Figure 3D. While previous ensemble modeling results indicated several enzyme overexpressions that might increase lysine pathway flux, our reimplementation demonstrates that the observed sequential overexpression experiments can be recreated through a wide variety of possible parameterizations, with a resulting wide distribution in possible flux responses. These results show that the method generalizes well to the case where insufficient data is provided to constrain model predictions and underscores the importance of rigorously characterizing posterior parameter space to determine the full range of possible model responses.

**Figure 3:**
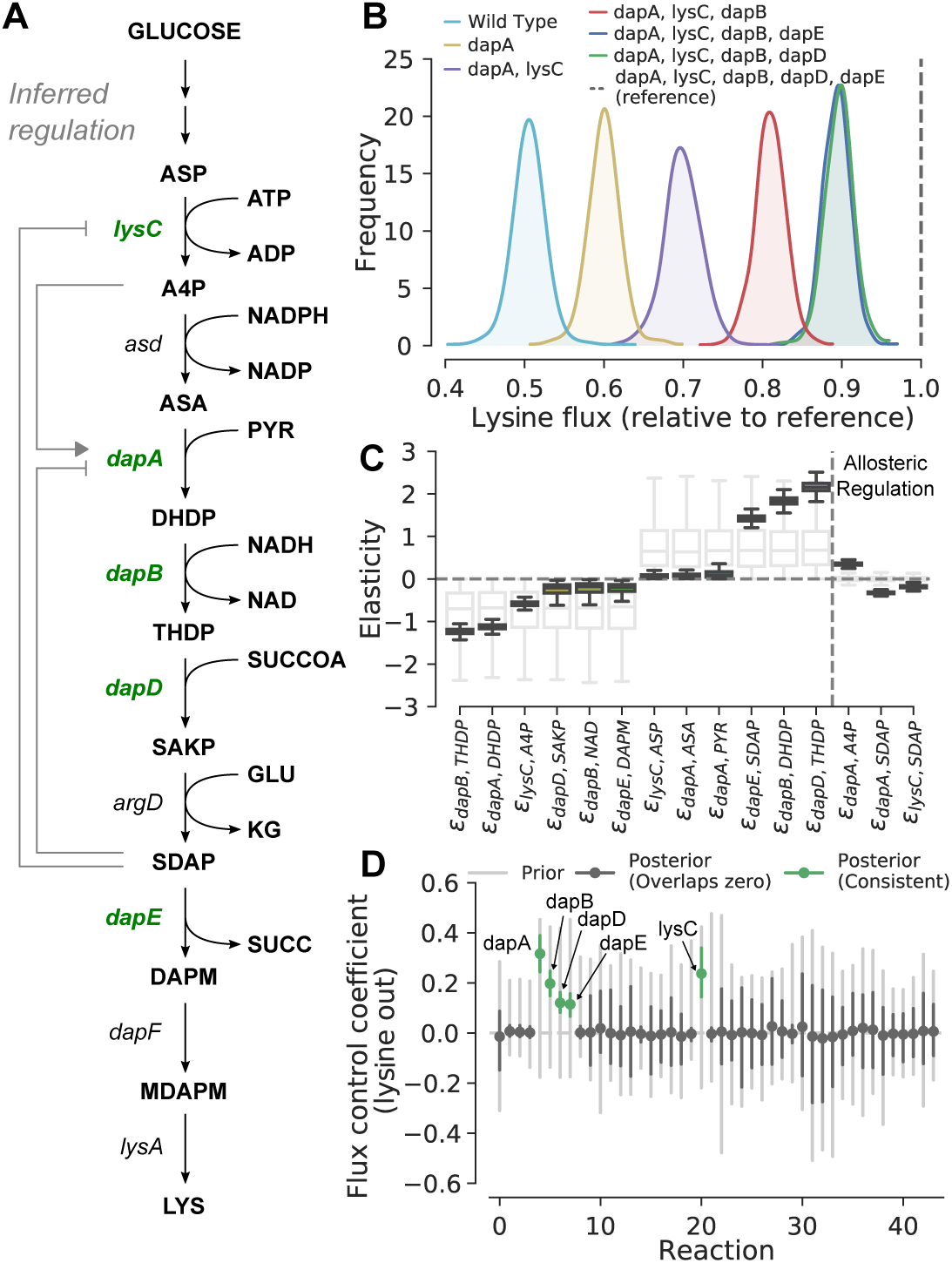
Inference on a medium-scale metabolic network with limited data. (A) Schematic of a portion of the considered metabolic network corresponding to lysine biosynthesis. Reactions shown in green were experimentally determined to improve lysine yields. Regulatory elasticities that were confidently inferred by the model are shown in gray. (B) Posterior predictive distributions for the enzyme overexpression experiments. Since the reference state was chosen as the highest-producing strain, all other strains have a relative lysine flux less than one. (C) Distributions of elasticities informed by the experimental results. Prior distributions for these elasticities are shown in light gray. Allosteric elasticities with Laplace priors that are confidently inferred are shown as the last four entries. (D) Flux control coefficients for each reaction in the model. Prior distributions (light gray) are mostly centered around zero. Posterior distributions (dark gray) are highlighted in green if their 95% HPD does not overlap zero. All lines indicate 95% HPD ranges, dots indicate median.

### Informing strain design through multiomics

The main strength of the proposed method is its ability to constrain kinetic parameters using multiomics data, even for large-scale metabolic systems. We therefore demonstrate the method using literature data on metabolomics, proteomics, and quantification of exchange fluxes for 25 different chemostat experiments with yeast (Hackett et al., 2016). The dataset comprises 5 different media conditions, each of which was run at 5 different dilution rates. We adapt a large-scale metabolic model of yeast metabolism that includes many of the genes, metabolites and boundary fluxes of interest (from (Jol et al., 2012)). The adapted model contains 203 metabolites and 240 reactions and was obtained by removing blocked metabolites and reactions under growth on glucose. As the goal in this example is to demonstrate that linlog kinetics are able to consume large amounts of multiomics data, a reference state near the center of the considered data was chosen, specifically the chemostat with phosphate-limiting media at a 0.11 hr^-1^ dilution rate. Reference fluxes (*v*^∗^) were calculated by minimizing error with the experimental boundary measurements while enforcing a nonzero flux through each reaction. In total, the experimental data consists of 1800 metabolite measurements, 792 boundary flux measurements, and 3480 enzyme measurements (omitting the reference state). Since the linlog inference framework only uses relative changes to enzyme, flux, and metabolite concentrations with respect to a reference state, it can naturally ingest large-scale multiomics datasets without the need for absolute quantification. In this example, relative metabolite concentrations are given as log2-transformed values (Boer et al., 2010). Even with an unknown pre-exponential constant *A,* relative concentrations *χ*can be calculated from log2-transformed concentrations *a* and *b:*

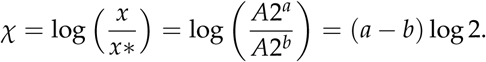

Distributions of the transformed data are shown in Figure 4A, indicating that the majority of data falls within one order of magnitude from the reference state value (values shown are natural logs).

In fitting the observed steady-state phenotypes, the model has to account for not only experimental error in measured enzyme concentrations, but also for potential changes in gene expression in unmeasured enzymes. Allowing all enzyme concentrations to vary induces a trade-off where steady-state fluxes are controlled through changes to enzyme expression instead of changes to steady-state metabolite concentrations. While Hackett et al., 2016 have previously shown that metabolic control is mainly determined by metabolite concentrations, some mechanism for adjusting enzyme levels is required to buffer against errors in model formulation and experimental measurements. We therefore place prior distributions on log enzyme concentrations for each condition that drive enzyme changes towards their measured values, or, if the reaction is not measured, towards zero (unchanged):

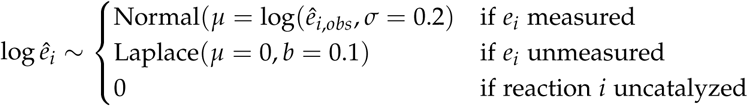

By placing a Laplace prior on unmeasured enzymes, we create a regularizing effect that penalizes an over-reliance on enzymatic control. Thus, we allow unmeasured enzyme concentrations to deviate from zero only if there is sufficient evidence. The model also has to consider changes in the external metabolite concentrations between media formulations and dilution rates. We therefore place vague priors on the external concentrations of imported substrates, including glucose, phosphate, sulfate, nitrogen, and oxygen:

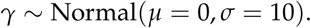

The model parameters therefore include 915 elasticities associated with direct kinetic regulation, 23,684 elasticities associated with potential off-target allosteric regulation, 4,680 enzyme expression levels (195 enzymes over 24 experiments), and 192 external metabolite concentrations (8 metabolites over 24 experiments), for a total of 29,471 parameters. While this number is far greater than the number of experimental data points, regularization forces many of these parameters to be zero.

Observed steady-state metabolite concentrations and fluxes are incorporated through a likelihood model that assumes experimental error is normally distributed around log-transformed metabolite and boundary flux data. Standard deviations were chosen as *σ_x_* = 0.2 for the metabolite data and *σ_v_* = 0.1 for the log-transformed fluxes. To improve numerical stability, we also clip the log-transformed, relative experimental data to ±1.5, such that log-transformed experimental data and model predictions greater than 1.5 or less than -1.5 are replaced by ±1.5. This process has the effect of reducing the influence of extreme points, especially in regimes far from the reference state that are unlikely to be fit well by the linlog approximation. However, the model is still required to predict the directionality and high-magnitude of these points correctly. Fitting the model using ADVI required 40,000 iterations of stochastic gradient descent, taking approximately five hours on a single compute node (Figure 8). The model is able to recapture a vast majority of the variance seen in the experimental fluxes, enzymes, and metabolites. Median absolute errors between the model predictions (median of the posterior predictive distribution) and experimental data points are 0.124, 0.0952, and 0.0186 for log-transformed metabolite, flux, and enzymes, respectively, for normalized points that fall within the [-1.5, 1.5] window.

**Figure 4:**
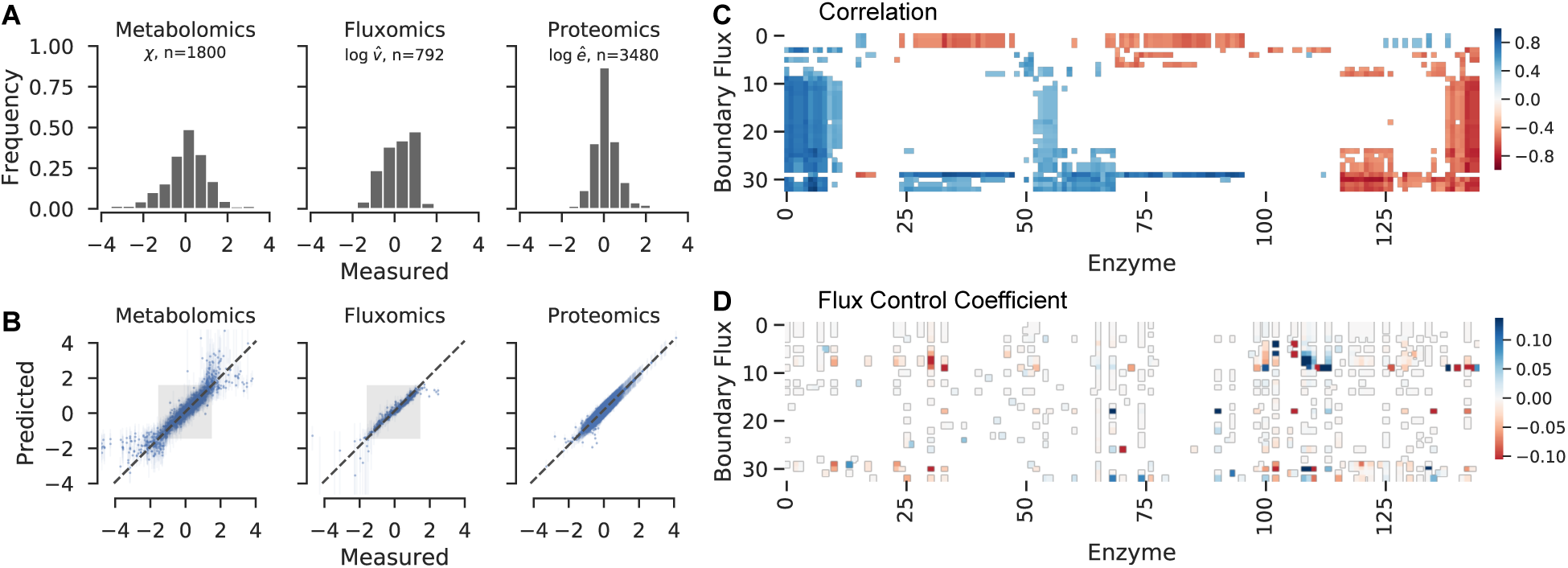
Parameterizing a genome-scale kinetic model with multiomics data. (A) Distributions in log-transformed experimental data after normalizing with respect to the phosphate-limited reference state. (B) Posterior predictive distributions after fitting with ADVI. Higher weight was given to experimental datapoints close to the reference state (±1.5) as indicated by the gray boxes. (C) Heat map of correlation coefficient between experimental enzyme measurements (x-axis) and experimental boundary flux measurements. Boundary fluxes and enzymes are sorted with hierarchical clustering. (D) Heat map of flux control coefficients as estimated from posterior parameter distributions. Boundary flux and enzyme ordering match those determined in (C). Colors represent medians of the posterior predictive distributions, FCCs with a direction that could not be confidently determined (having a 95% HPD that crossed zero) are colored white.

Posterior distributions in fitted parameter values indicate that the model is able to fit the observed experimental data while using relatively few of the additional regulatory parameters. Of the 23,684 regulatory elasticities, only 153 (0.65%) were confidently nonzero. However, we note that determining mechanistically accurate regulatory interactions from observations of steady-state flux behavior is inherently difficult. For instance, for a regulatory pathway in which A regulates B and B regulates C, identifiability issues might cause the pathway to be modeled as A regulates B and A directly regulates C. While poorly identifiable, the impacts of these alternative regulatory topologies on flux control coefficients are largely similar. Of the 50 unmeasured enzymes, only half were nonzero in at least one experimental condition. Overall, only 35% of the available unmeasured enzyme expressions differed from their reference state value.

We next look at what the model is able to learn about the systems-level control of yeast metabolism. A common goal in strain engineering is to find gene targets for increasing the yield of a given metabolic product. We therefore look at relationships between enzymes with measured protein concentrations and measured boundary fluxes. In a traditional statistical approach, correlations between enzyme levels and metabolite fluxes might be used to further enhance production of a desired metabolite. Figure 4C shows a heat map of Pearson correlation coefficients between enzyme expression levels (as determined through proteomics) and measured metabolite boundary fluxes. A permutation test was performed to determine correlations significant at the *α* = 0.05 confidence level; non-significant correlations were masked from the array. In this map, hierarchical clustering is used to reveal clear groups of metabolites and enzymes that vary together in the experimental data. A larger version of this image, with labelled axes, is shown in Figure 9. However, correlations between proteins and metabolite boundary fluxes do not necessarily imply that a particular enzyme is involved in directing flux to a particular product. For instance, several of the highest correlations exist between methionine synthase and relatively distant amino acid products alanine, arginine, and histidine. The top ten enzyme-boundary flux correlations are shown in Table 2.

**Table 2:**
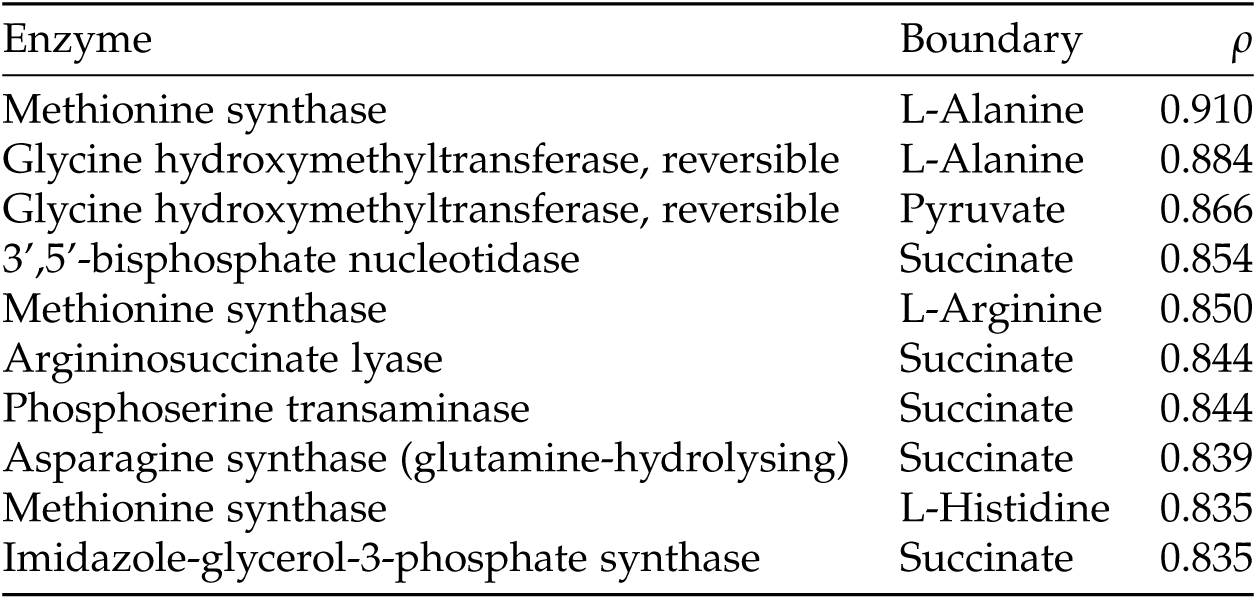
Largest significant correlations between measured enzymes and measured boundary fluxes.

Flux control coefficients as estimated through the proposed method therefore offer an alternative approach for determining potential enzyme targets that more systematically considers the effects of metabolic stoichiometry and kinetics. Before considering posterior distributions in flux control coefficients, we first look at whether the prior assumptions on enzyme elasticities and model stoichiometry esult in any confidently nonzero values. From the prior predictive distribution, only 6 enzyme-boundary flux pairs have a significantly nonzero FCC, and typically involve reactions directly associated with metabolite production. For instance, a positive flux control coefficient is associated with asparagine synthase and valine transaminase on asparagine and valine export, respectively. A heat map of FCCs calculated from the fitted posterior elasticity matrix is shown in Figure 4C, in which FCCs that have a 95% HPD that includes zero are colored white. Unlike the map of correlation coefficients, FCCs result in a much sparser matrix of inferred connections between enzyme concentration and steady-state flux. However, these coefficients are much more interpretable as direct causality between enzyme expression and increased downstream flux. The top 10 largest, identifiable flux control coefficients are shown in Table 3. Some pairs of enzymes and boundary fluxes, *i.e*. glycerol-3-phosphate dehydrogenase enhancing glycerol production, are direct upstream enzymes for the boundary flux in question. However, since linear pathways can have an uneven distribution of flux control coefficients, determining the rate-limiting step in biosynthesis pathways is an important result. Other confident FCCs represent more indirect effects, for instance the consumption of the upstream phosphoenolpyruvate in 3-phosphoshikimate 1-carboxyvinyltransferase reducing the export of pyruvate.

**Table 3:**
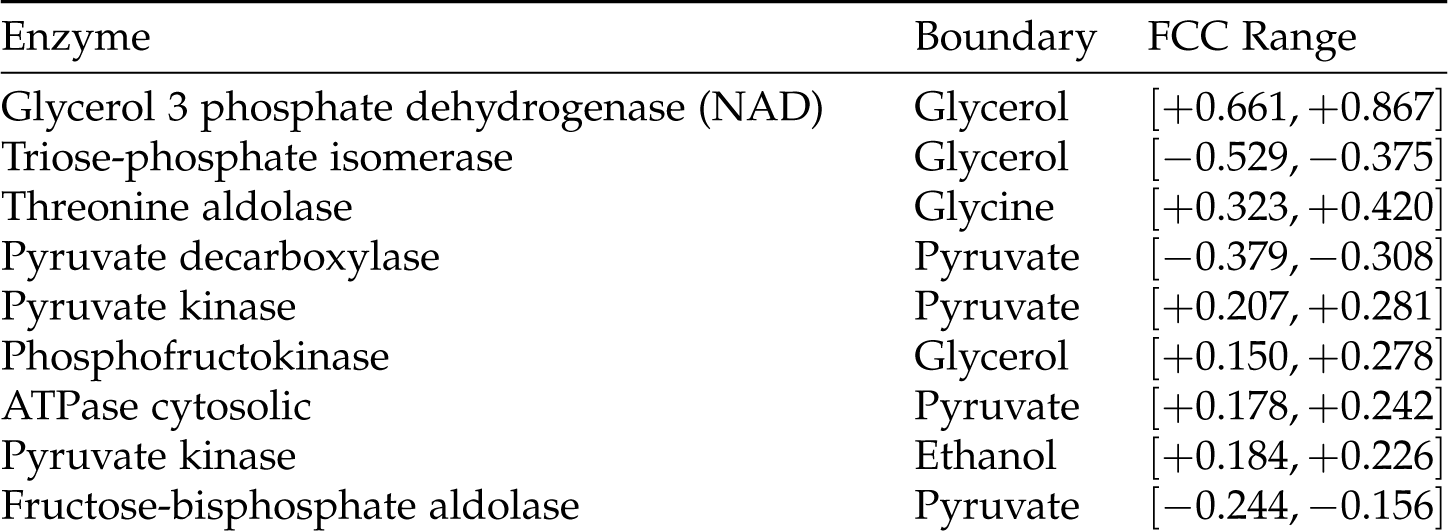
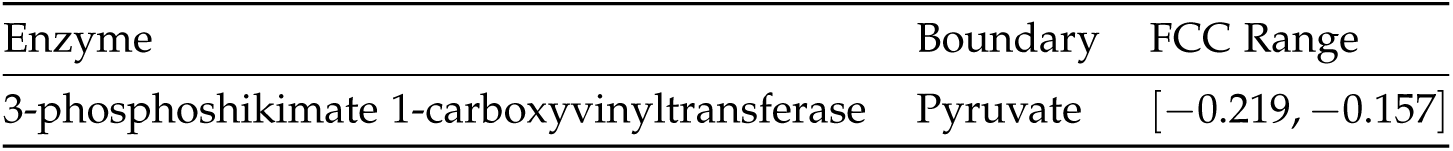
Largest flux control coefficients for the modulation of measured enzymes on measured boundary fluxes. FCC ranges represent upper and lower bounds of the 95% highest posterior density. Enzyme-boundary pairs that also appear as confident predictions prior to including experimental data are omitted.

## Methods

All simulations were performed in Python using the pymc3 library (Salvatier et al., 2016). Additional code to initialize the elasticity prior matrices and calculate the steady-state metabolites and fluxes is provided at github.com/pstjohn/emll, along with jupyter notebooks detailing the use cases described above.

## Conclusion

In this study we demonstrate how kinetic models of microbial metabolism can be analyzed through modern probabilistic programming frameworks. In doing so, we have invoked approximate formalisms for enzymatic kinetics; however, we note that similar trade-offs between modeling fidelity and computational efficiency are common throughout biology and chemistry. For instance, while small-scale pathways might be better modeled at a higher level of kinetic theory, a complete kinetic description of a genome-scale kinetic model is likely currently infeasible given available data and computational resources. As biological experiments are becoming increasingly easy to iterate with modeling results, a complete kinetic description of a given pathway may not be as valuable as a reasonable guess as to how to improve a desired phenotype. Computational methodologies that quickly converge to generate a list of potential targets, such as the one proposed in this study, may therefore be essential in keeping up with the growing ease of multiomics experiments. The proposed method can also be run efficiently on consumer-grade hardware, a important factor for applications in industrial microbiology where access to large-scale high performance computing resources is limited.

As the field of variational inference is rapidly evolving, this technique could likely be made more robust or efficient through the use of alternative inference algorithms. For instance, correlations between elasticities were demonstrated through a Hamiltonian Monte Carlo trace but were missed by the corresponding mean-field Gaussian approximation. While fitting a full-rank Gaussian is likely impractical at larger data set sizes, reduced-rank approximations (Rezende and Mohamed, 2015) might offer a suitable compromise between posterior accuracy and computational efficiency. Additionally, inference approaches which only consider a subset of the experimental data might also prove useful. Since each perturbed state involves a new linear solve in calculating the likelihood, stochastic variational inference (Hoffman et al., 2013) or firefly MCMC (Maclaurin and Adams, 2014) might reduce the cost of approximating or drawing samples from the posterior.

## Acknowledgement

This work was authored in part by Alliance for Sustainable Energy, LLC, the manager and operator of the National Renewable Energy Laboratory for the U.S. Department of Energy (DOE) under Contract No. DE-AC36-08GO28308. Funding was provided by the U.S. Department of Energy Office of Energy Efficiency and Renewable Energy Bioenergy Technologies Office via the Agile BioFoundry to PCSJ. We thank Jay Fitzgerald at DOE, Jacob Hinkle, and members of the Agile BioFoundry for helpful discussions.

## Supplementary Material

**Figure 5:**
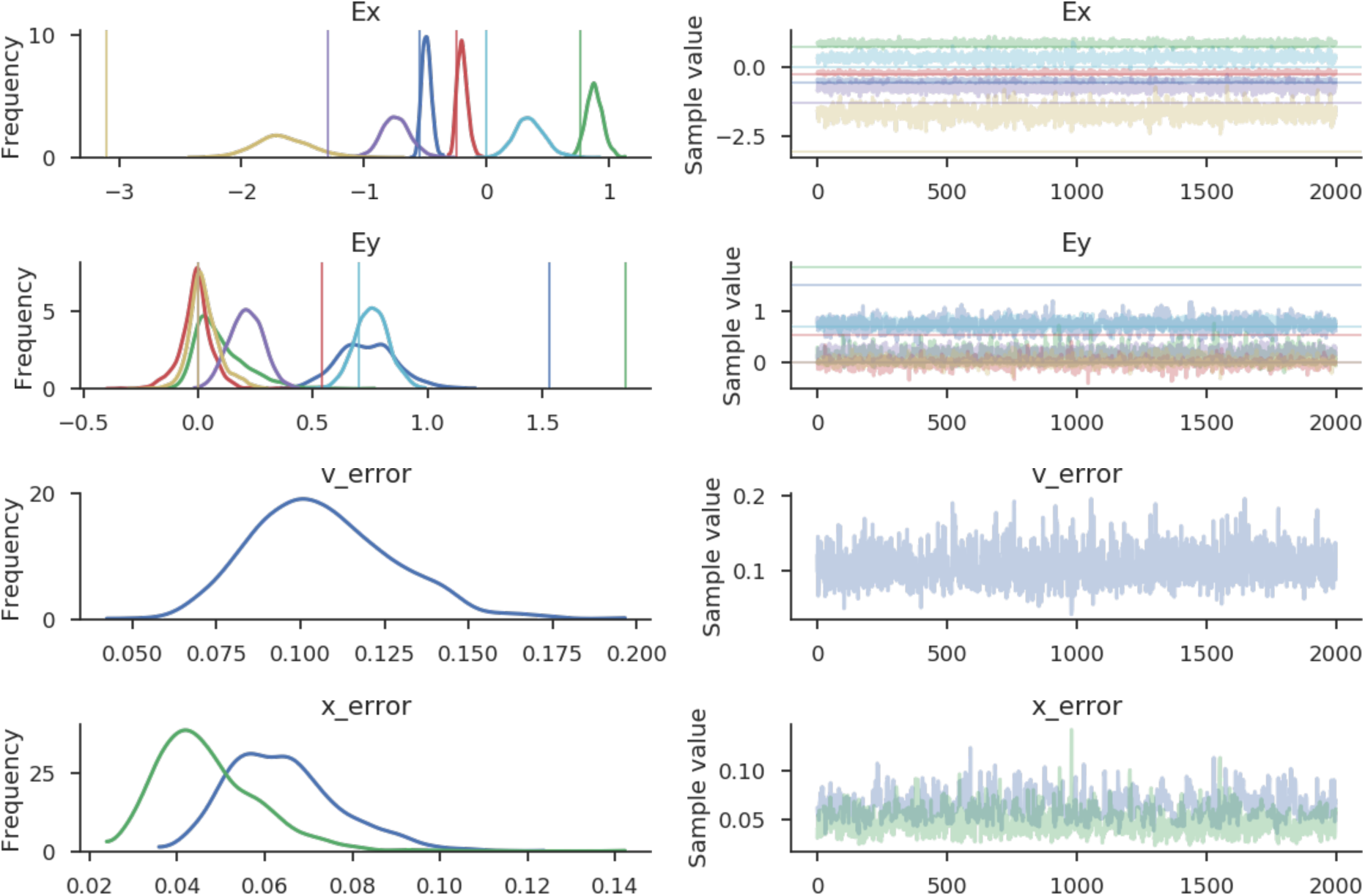
Trace of the NUTS sampler for the *in vitro* dataset. (left) kernel density estimates of each parameter. Vertical bars indicate the values obtained using the multiple linear regression technique of (Wu et al., 2004). (right) Samples from the MCMC sampler

**Figure 6:**
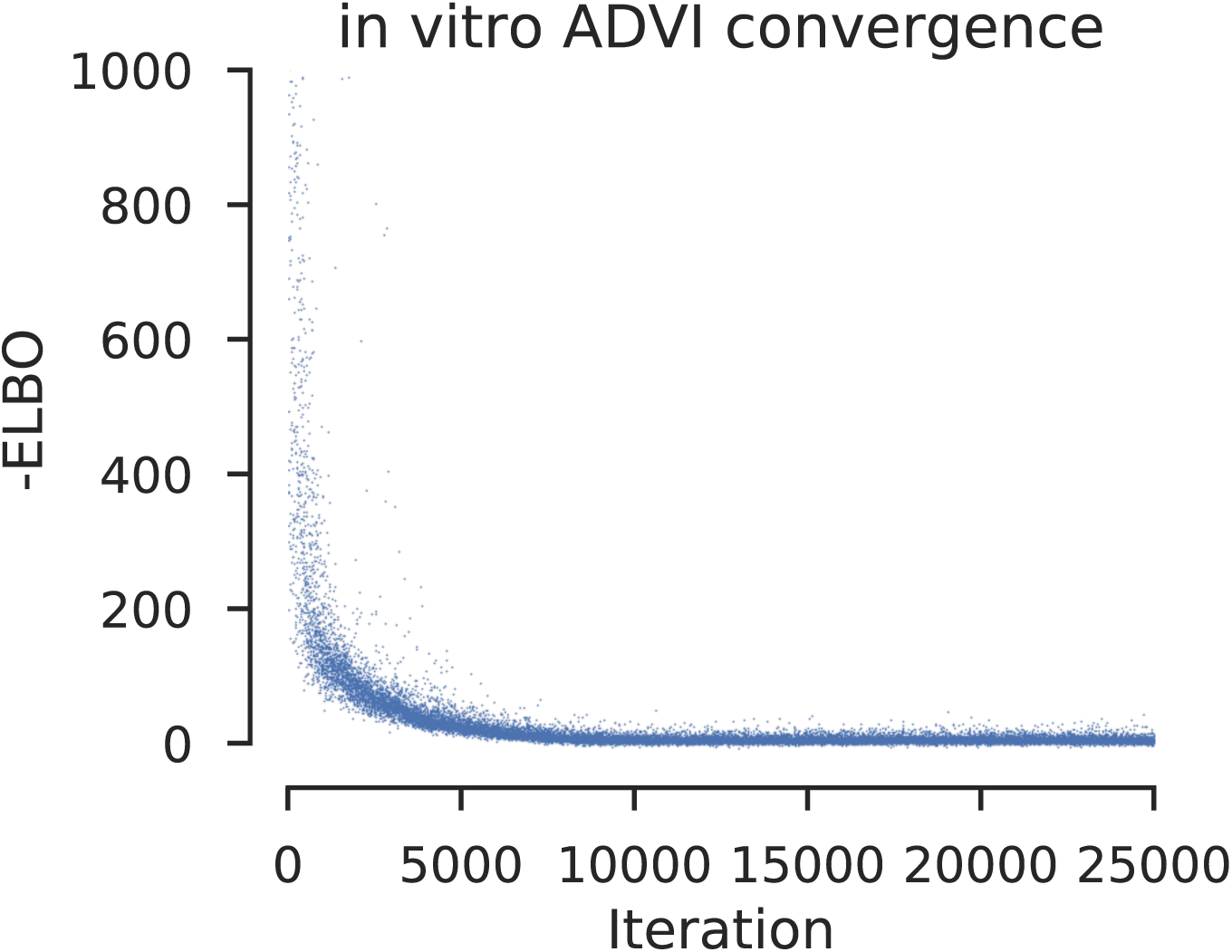
Convergence of the Evidence Lower Bound (ELBO) for the *in vitro* dataset.

**Figure 7:**
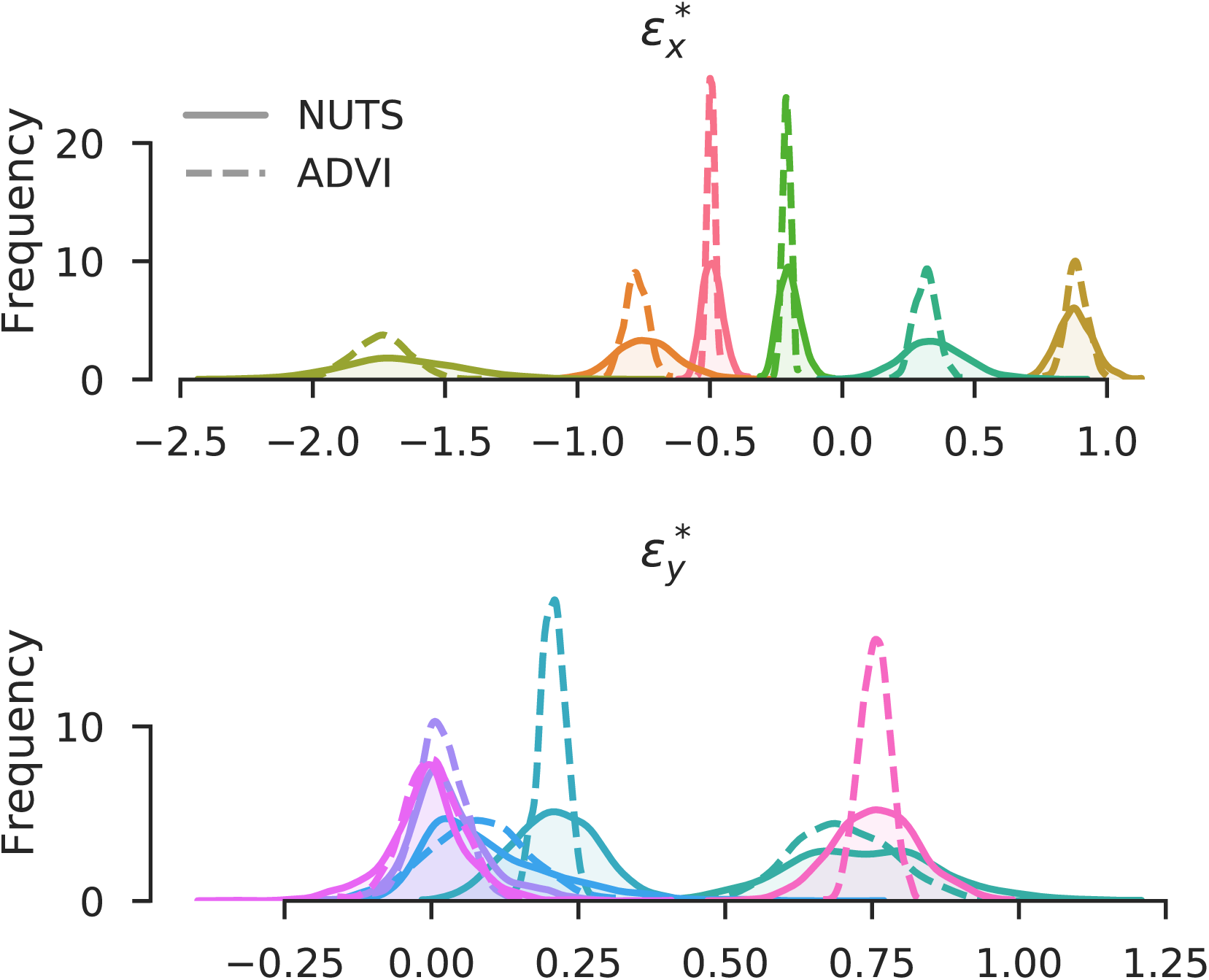
Comparison between posterior distributions for the *in vitro* dataset as estimated by NUTS (solid lines) or ADVI (dashed lines). ADVI posteriors have a similar mean but smaller variance.

**Figure 8:**
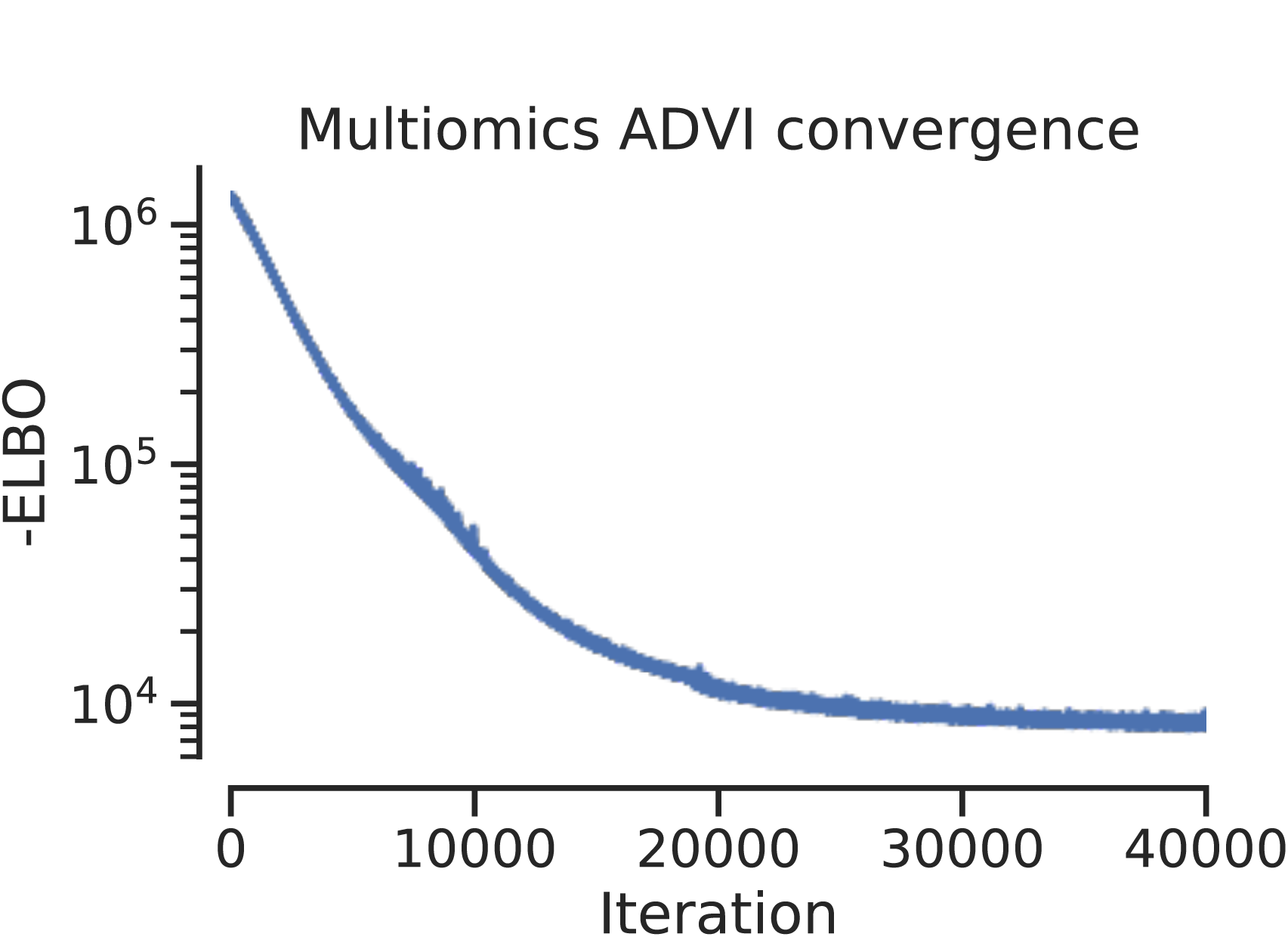
Convergence of the Evidence Lower Bound (ELBO) for the multiomics dataset and yeast metabolic model

**Figure 9:**
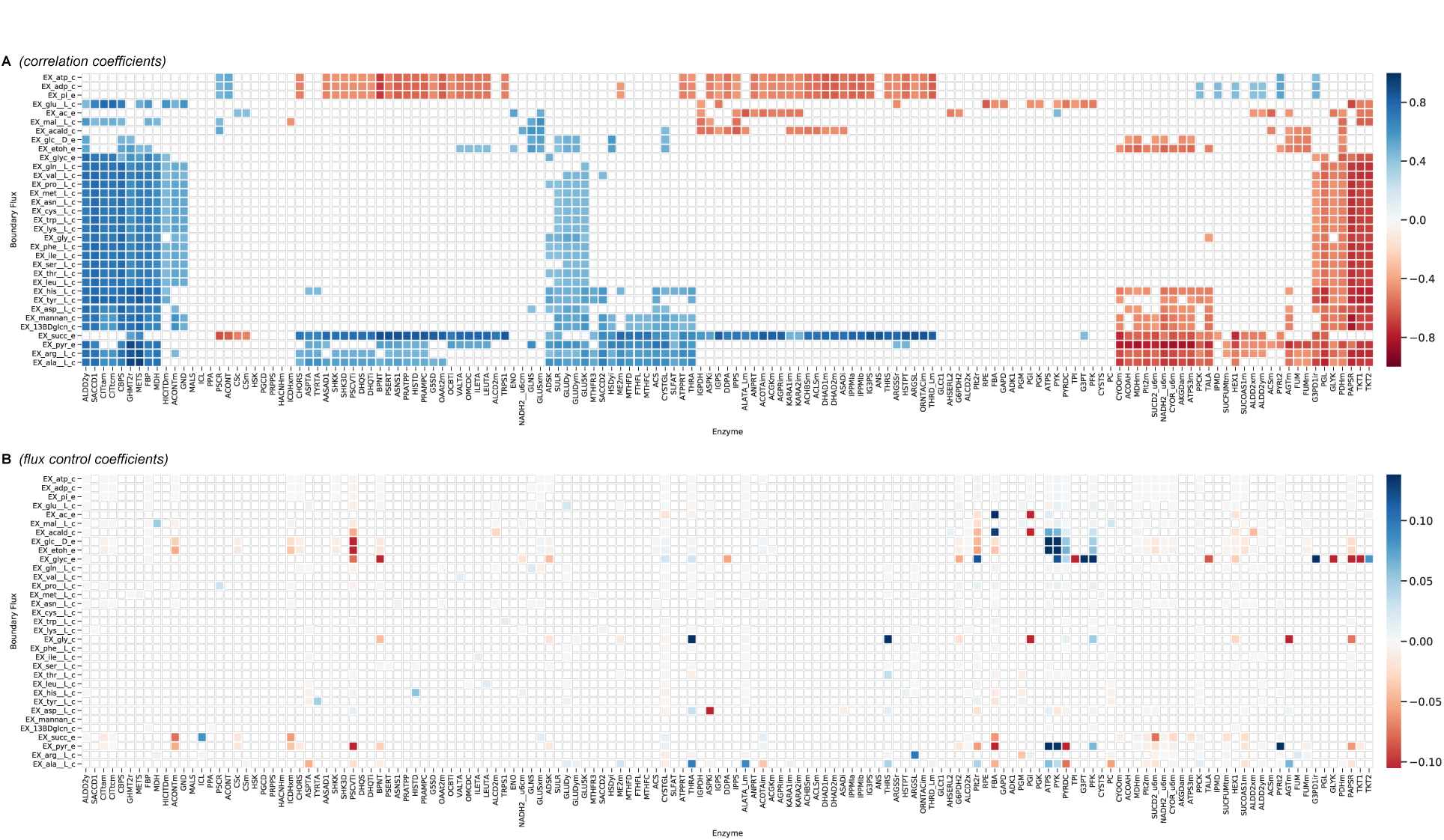
Full heatmaps (with labeled boundary fluxes and enzymes) for the multiomics dataset. Full names for the reaction IDs shown can be found in the detailed model description.

### Calculating reverse-mode gradients for regularized linear solve

In order to efficiently perform many inference approaches, forward and reverse-mode gradients for the likelihood function are required. In this method, the least-squares linear solve is a particularly tricky operation for which gradients in some automatic differentiation packages are not automatically supplied. In this section, we therefore derive the necessary matrix equations to calculate forward and reverse mode gradients for the least-norm linear solve, *χ_ss_* = **A**^†^**b**. In practice, it is much more efficient to calculate this least-norm solution directly *(i.e.,* using the LAPACK routine dgelsy) instead of explicitly calculating the pseudoinverse matrix.

Gradients for the least-norm solution are derived by first calculating those for Tikhonov regularization, and subsequently taking the limit as *Λ* → 0. Definitions for matrix derivatives are taken from (Giles, 2008). Similar to example 2.3.1 in Giles (2008), the forward derivative for a Tikhonov-regularized linear takes the form

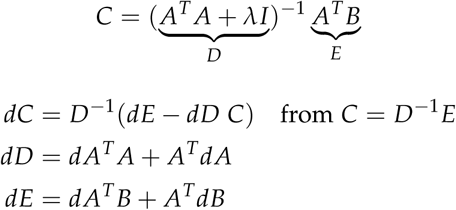

Substituting into the equation for *dC,*

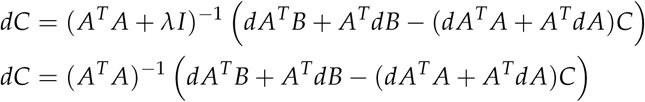

Also following Giles (2.3.1), the reverse mode gradient can be found:

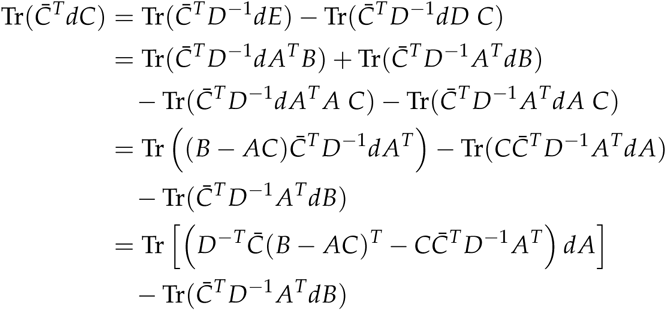

therefore,

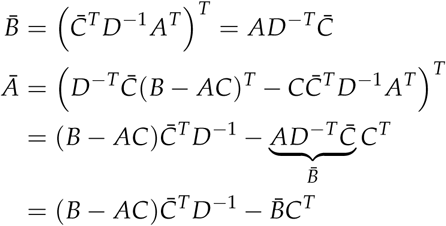

Since *D* = *A^T^ A* + *λI* = (*A^T^ A* + *λI*), these gradients can be further simplified using the relations

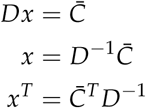

After substituting into the equations for 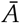 and *B,* we are left with

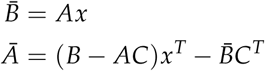

as the reverse-mode gradients for the least-norm solve *C* = *A*†*B*.

